# Icariin prevents depression-like behaviors in chronic unpredictable mild stress-induced rats through Bax/cytoplasm C/caspase-3 axis to alleviate neuronal apoptosis

**DOI:** 10.1101/2023.03.09.531962

**Authors:** Xiao Wu, Xiaona Zhang, Lulu Sun, Xiaomin Lu, Cunsi Shen

## Abstract

Major depressive disorder (MDD) affects approximately 16% of the global population. Our previous study has demonstrated that icariin (ICA) exhibits anti-depressant activity by increasing the expression of Brain Derived Neurotrophic Factor (BDNF) in a rat model of chronic unpredictable mild stress (CUMS). In this study, we investigated whether and how ICA can prevent CUMS-induced depression-like behaviors in rats by modulating hippocampus neuronal apoptosis. Forty male rats were randomly divided into control, CUMS, CUMS-fluoxetine (Flx) (10 mg/kg), and CUMS-ICA (20 mg/kg) groups. Behavior tests including sucrose preference test (SPT), open field test (OFT), elevated plus-maze (EPM), and forced swimming tests (FST) were performed. The Nissl staining and TUNNEL assay were used to determine neuronal apoptosis. Subsequently, expression of glucocorticoid receptor (GR), Bcl-2, cytochrome C, caspase-3 and Bax in the hippocampus were tested by western blot. Our results show that a chronic administration of ICA (20 mg/kg) can prevent CUMS-induced depressant-like behaviors in male model rats. Additionally, ICA significantly inhibited mitochondrial translocation of GR, reduced mitochondrial outer membrane permeabilization (MOMP) to suppress the release of cytochrome C, and then inhibit the activation of caspase-3. In conclusion, our research provides new evidence to understand the anti-depressant activity of ICA, which relates to its inhibition of neuronal apoptosis in hippocampus through mitochondrial apoptotic pathway.

## Introduction

Stress, especially psychosocial stress, plays a crucial role in the pathogenesis of major depressive disorder (MDD) [1]. Now, mental disorders account for a large proportion of the burden of disease in governments around the world, surpassing cardiovascular and cancer diseases [2]. According to the World Health Organization, depression will become the second leading cause of disability in 2030. Although several antidepressants targeting the serotonin and/or norepinephrine systems have been used to treat depression [3, 4]. There is still no evidence of a reduction in the population burden of depression. One possible explanation is that treatment may not be adequately available, effectively available to have an impact [5]. So, different strategies, such as preventive or alternative medicine, need to be further explored [6].

The hippocampus is a vital component of brain, it participates in several important functions, including behaviors, mental and intellectual activities, in both rodents and human [7]. Morphological changes in brain tissues are observed with long-term MDD, in particular a decreased volume and neuronal apoptosis of the hippocampus [8]. Physiologically, when the hypothalamus-pituitary-adrenal (HPA) axis is activated, the adrenal glands secrete glucocorticoids (GCs). During chronic stress, dysfunction of the HPA axis is accompanied by significant changes in neuroendocrine function. HPA axis activation can be regulated by negative feedback mechanisms that activate glucocorticoid receptor (GR) at different locations, including the hippocampus, prefrontal cortex (PFC) and upper brain structures [11], where dissociated GR signals to the nucleus and regulates the target genes. Mitochondria are essential organelles that regulate cellular homeostasis and cell survival [12]. More and more evidence suggests that MDD may be a consequence of abnormal mitochondrial function [13]. Due to a vital role of in cell physiology, mitochondria should be the first responder to stress. Animal studies have also shown that CUMS inhibits mitochondrial oxidative phosphorylation, dissipates mitochondrial membrane potential, and disrupts the mitochondrial ultrastructure of various brain regions, including mouse hippocampus, cortex and hypothalamus [14]. For example, Rudranil De et al. revealed that Bax can act as the central mediator by translocating into mitochondria and inducing neuronal apoptosis when brain tissue is stimulated by stress [15]. In addition, mitochondrial transcription and energy metabolism are also affected by GCs [16]. The effect of GCs on mitochondrial processes can be explained by GR translocation to mitochondria [17]. The previously confirmed mitochondrial translocation of activated GR in DP thymocytes provides an intriguing explanation for its marked sensitivity to GC-induced apoptosis [18]. However, the fine molecular details of how the mitochondrial translocation of GR regulates neuronal apoptosis remained unclear.

Icariin (ICA) is a flavonoid glucoside isolated from *Epimedium brevicornu Maxim*, which is frequently used in traditional Chinese medicine (TCM) to treat asthma [19], kidney disease [20], testicular dysfunction [21] and cartilage damage [22]. TCM is often used to treat depression, for example, paeoniflorin improves chronic stress-induced depression behavior in mice model by influencing the ERK1/2 pathway [23]. Zhang et al. revealed that Xiaoyao powder can reduce the damage of hippocampal neurons in CUMS-induced hippocampal depression model rats through connexin 43Cx43/GR/brain-derived neurotrophic factor signaling pathway [24]. Classically, ICA has also been reported to exert antidepressant-like actions. Our previous work showed that ICA improves hippocampal neuroinflammation by inhibiting HMGB1-associated pro-inflammatory signals in LPS-induced inflammatory models of C57BL/6 J mice [25]. In addition, ICA by inhibiting NF-κB signal activation and NLRP3-inflammasome/caspase-1/IL-1β axis plays an antidepressant-like role in CUMS model of depression in rats [26]. Otherwise, ICA reduces Glu-induced excitatory neurotoxicity through antioxidant and anti-apoptotic pathways in SH-SY5Y cells [27]. However, whether ICA is beneficial for the depression via ameliorating neuronal apoptosis in the hippocampus is still unknown.

In this study, the CUMS protocol was used as a rat model for depression, which mimicked many symptoms of human depression [34]. Understanding the molecular regulatory mechanism of neuronal apoptosis under the CUMS-induced depression-like behavior in rat model may provide the novel therapeutic targets for depression. Therefore, the purpose of the present study was to identify the anti-apoptosis mechanism of ICA in the hippocampus on the CUMS-induced depression-like behavior in male rat model. In addition, GR, Bax, Bcl-2, Caspase 3, Cleaved Caspase 3, and cytochrome C levels in the hippocampus were measured to explain the possible mechanism of ICA. Here, we provided evidence for the activation of the mitochondrial apoptotic pathway associated with GR after ICA treatment in hippocampus, which support ICA might act as an important drug in the prevention of depression.

## Methods and materials

### Drugs and reagents

ICA (the purity is 98.93%) was purchased from Shanghai Ronghe Medical Science Co., Ltd. (Shanghai, China). Dimethyl sulfoxide (DMSO) was used to prepare ICA stock solutions and diluted with sterile normal saline (DMSO concentration: 0.1%) [35]. Fluoxetine (Flx) was purchased from Eli Lilly and Company (Suzhou, China) and diluted to 10 mg/mL with sterile saline solution [36]. Mitochondria Isolation Kit for Tissue (No.89801) was bought from Thermo Fisher Scientific Inc. Rat corticosterone (CORT) ELISA kits from ebioscience (San Diego, CA) were purchased from Beyotime Biotech Inc., China. Mouse anti-GR (1:1000), mouse anti-Bcl-2, mouse anti-Bax (1:400, Santa Cruz Biotechnology, Santa Cruz, CA, USA), anti-β-actin, anti-Cox-IV, anti-caspase-3, cleaved caspase-3, anti-cytochrome C (1:1000, Bioworld Technology Co., Ltd, Nanjing, China). Tunnel was bought from Wuhan Boster Co., Ltd. (China).

### Animals

All experimental male Sprague-Dawley (SD) rats weighing 120-140 g (5 weeks old) were purchased from Shanghai SLAC Co. (Shanghai, China). The animals were kept in temperature (20 ± 2 °C), humidity (50–60%) and 12 h light/dark cycle (light period from 6:00 am to 18:00 pm) and free of food and water *ad libitum*. To avoid fighting among male rats, prior to the experiment, all animals were maintained in a single cage for at least seven days [37]. All experiments were conducted in accordance with the guidelines of the Animal Care and Use Committee at Huashan Hospital of Fudan University. Make every effort to reduce the suffering of animals. In addition, all procedures were approved by the Animal Care Ethics Committee of Huashan Hospital of Fudan University (approval number: [HS-A-2021-0721]).

### Chronic Unpredictable Mild Stress (CUMS) procedure and drug treatments

Forty rats were randomly divided into 4 groups (10 rats/group): Control, CUMS-vehicle, CUMS-Flx (positive control, 10 mg/kg), and CUMS-ICA (20 mg/kg). Control group rats were grouped housed in a separate room under standard conditions. All rats that underwent the CUMS procedure were single housed. The CUMS procedure was performed in one of the two rooms according to our previous protocol [38]. The dose of ICA was chosen on the basis of our previous experiments in rats showing that ICA had a significant impact on behavior at 20 mg/kg [25].

The three groups exposed to CUMS underwent the sequence of stressors for 5 weeks, and were administered with vehicle (saline 10 ml/kg), ICA (20 mg/kg), or Flx (10 mg/kg) by gavage at 11:00 a.m, respectively, once daily for the 35 days of CUMS. After 5 weeks of CUMS exposure, rats were subjected to different behavioral tests at least 16-18 h after the last dose. Time schedule and CUMS procedure of experiments are illustrated in Figure 1.

**Fig.1.**
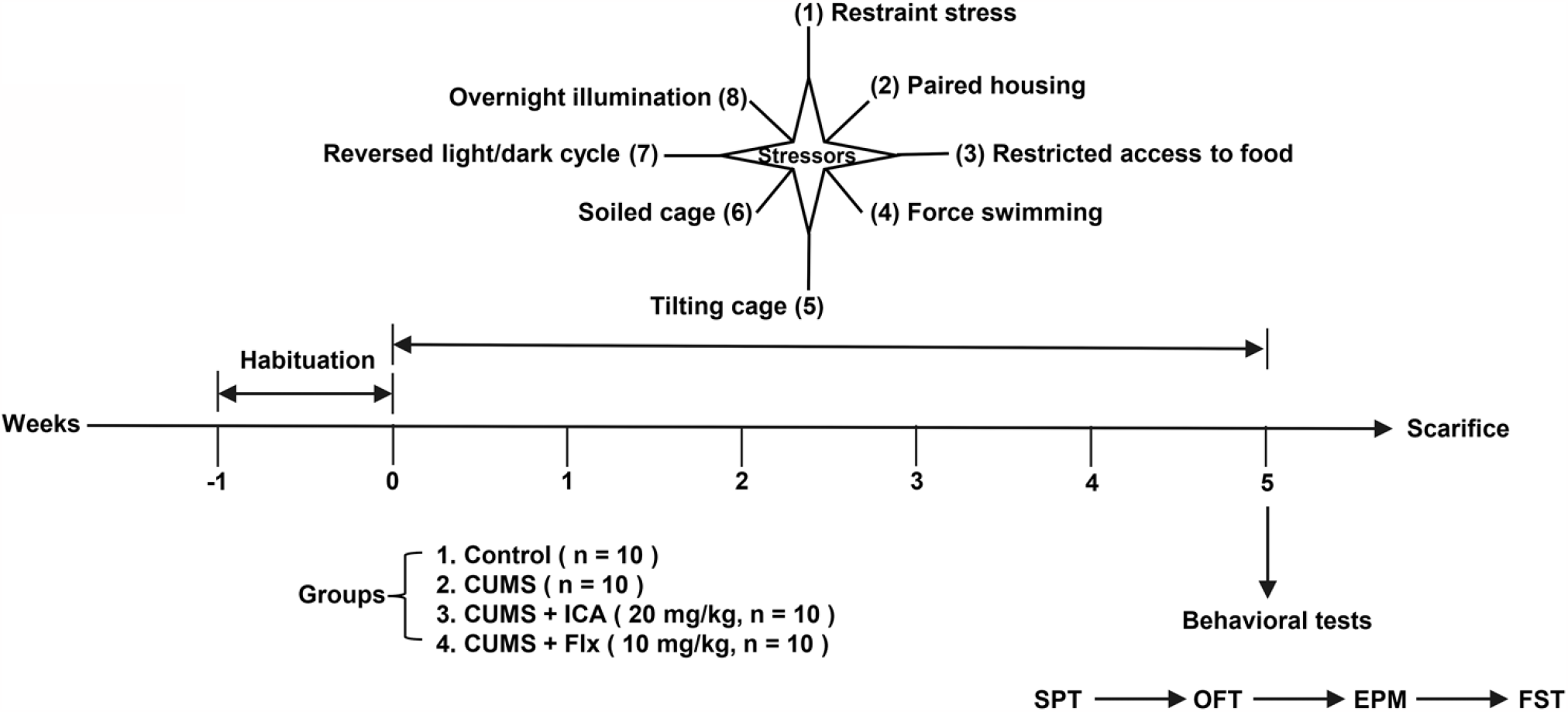
Timeline of the experiment. Before the experiment, the animals were allowed a 7-days adaptation period. Except for the control group, the rats in the other groups were subjected to CUMS for 5 weeks and treated with different drugs. CUMS procedure was performed in random order. The body weights were monitored weekly. SPT, OFT, EPM and FST were performed in order at day 35. Then, the rats were sacrificed for further detection.

### Behavioral Tests

Before the experiment, the animals were allowed one week adaptation period. All experiments were conducted between 8:00 am and 14:00 pm on the 35th day to minimize the impact of circadian rhythm after the last treatment. In addition, rats were evaluated after 15 min of testing room habituation. Sucrose Preference Test (SPT) was performed first, followed by Open field test (OFT), Elevated Plus-Maze (EPM), and finally forced swimming test (FST) [39]. Double blindness was used for behavioral testing, and all animals underwent behavioral tests. Experiments were performed and repeated 3 times.

### Sucrose Preference Test (SPT)

The SPT is commonly used to quantify anhedonia was performed after five weeks and before surgery. Briefly, all rats were first adapted to two bottles of 1% sucrose solution for 24 h. For the next 24 h, rats were free to choose between a bottle of sterile water and a bottle of 1% sucrose solution. Prior to the test, the rats were deprived of food and water for 23 h. Rats were then presented with two pre-weighted bottles of sterile water and 1% sucrose solution. After 1 h, consumption of sucrose and water intake was measured. Sucrose preference was calculated by the following formula: sucrose preference = sucrose consumption/[water consumption+ sucrose consumption] × 100% [40, 41].

### Elevated Plus-Maze (EPM)

The procedure was performed as described previously [42]. Briefly, the EPM is 50cm high, which has two enclosed (50 cm *10 cm *40 cm) and open (50 cm *10 cm) arms, the open central area is 10cm*10 cm area. Onset of the experiment, rats were placed in the central area, faced one of the enclosed arms. The experiment was recorded on a video recording system in the next room for five minutes. The video system records the frequency and time the rats entered the Open arms during the testing time. (RD1108-EPM-R, Shanghai Mobile Datum Corporation, Shanghai, China).

### Open Field Test (OFT)

The OFT was conducted as described by Yan et al [43]. The open field is a 100 × 100×40 cm cuboid box with the black odorless plastic floor divided into 25 squares. The surrounding 16 squares were considered as the peripheral zone, while the remaining 9 squares were regarded as central zone (regards as the social area). The movements of rats in the peripheral zone were defined as protected behavior and the movements in the central zone were defined as exploratory behavior. Rats were putted in the central zone and their movements were recorded for 5 minutes with a video system. The number of entries in the central zone, the total time expended in the central zone and defecation were scored.

### Forced Swim Test (FST)

The FST described by Porsolt et al [44] was lightly alteration [40]. In brief, each rat was placed individually in a cylindrical plexiglass container (diameter: 18 cm, height: 50 cm) with 7-8 liters of water at 23±1 °C. The rats were put in the container for a 15 min training; 24 hours later, the rats were put in the container again for another 5 mins test. The immobility time during the test was recorded by two observers blinded to the experiment.

### Serum CORT assay

After the FST procedure, blood samples were collected individually by intracardiac puncture between 11:00 am and 13:00 pm to avoid fluctuations in hormone levels. And separated in a refrigerated centrifuge at 3,000 rpm for 15 min at 4 °C. Serum was stored at −80 °C till the assays. Tested the CORT concentration with ELISA kit. The OD value at 405 nm was measured on an ELISA plate reader. CORT concentration was quantitatively determined by comparing with the standard curve. Detection threshold = 150 pg/mL, coefficient of variation limit =9.6%, and concentration expressed in pg/mL [45].

### Nissl Staining

After the behavior testing, the rats were sacrificed immediately with anesthesia and perfused by trans-cardiac with 4% para-formaldehyde in 0.1 M phosphate buffer. The brain tissues were dissected and immersed the same concentration of formaldehyde. The brain tissues contained hippocampus were embedded with paraffin and cut into 8 μm thick serial sections. The sections were stained with Cresyl Violet and mages were taken at ×200 magnification with a microscope (Olympus AX-70) [39]. We examined the morphology of neurons in the dorsal hippocampus of both hemispheres.

### In situ labeling of DNA fragmentation

Apoptotic cells were detected by TUNNEL assay. 8 μm thickness coronal sections were putted in 1x terminal deoxynucleotidyl transferase buffer (Invitrogen, Carlsbad, CA) for 30 min, then, they reacted with terminal deoxynucleotidyl transferase enzyme (Invitrogen) and biotinylated 16-dUTP (Roche Diagnostics, Indianapolis, IN) at 37 °C for 60 min. The slices were washed with 2x SSC (150 mol/liter sodium chloride and 15 mol/liter sodium citrate, pH 7.4) for 15 min, and then washed with PBS for 15 mins two times again. Counterstaining nuclei with methyl green solution using avidin-biotin technique [46].

### Mitochondrial Fractionation

Fresh hippocampal tissue was weighed (100 mg) and washed with pre-cooled PBS and cut into fragments, then the hippocampal debris were homogenized with a Dounce homogenizer on ice at 700 μL bovine serum albumin (BSA)/Reagent A Solution. Mitochondria Isolation Reagent C (700 μL) was added to the tube and centrifuged at 700 × *g* for 10 min at 4 °C to remove nuclei and unbroken cells. Transferred the supernatant into a new tube and centrifuged at 11,000 × *g* for 15 min at 4 °C, the sediments we got were just the mitochondria. And the supernatant was then transferred into another tube and centrifuged at 12,000 × g for 10 min at 4 °C to get the cytoplasmic without mitochondria [47]. Both the pellet (mitochondria fraction) and the supernatant (cytoplasmic fraction) were stored for further testing. Cytoplasmic and mitochondria fractions purity was confirmed by incubating specific antibodies against β-tubulin (T9026, Sigma) and mHsp70 (MA3-028, Affinity Bioreagents) for each compartment, respectively [48]. Representative blots demonstrating the purity of compartments are presented in Figure S1. All samples exhibited proper separation, and no sample separation was unclear.

### Western blots

Extracted the cytoplasm and mitochondria protein from the hippocampus, quantified the protein concentrations with BCA (Beyotime, China). Loading buffer (0.1 MTris-HCl buffer (pH 6.8) containing 0.2 M DTT, 4% SDS, 20% glycerol and 0.1% bromophenol blue) was used to dissolve 40-60 μg equal volume protein samples. The samples were separated on 10% SDS-PAGE and then electrically transferred to PVDF membrane at 90 V. PVDF membranes were incubated with TBST (containing 5% skimmed milk) for 1 h at 37 °C and with primary antibodies at 4 °C for 24 h. The primary antibodies used were as follows: mouse anti-Bax, mouse anti-GR (1:1000), mouse anti-bcl-2, (1:400, SantaCruz Biotechnology, USA), anti-β-actin, anti-Cox-IV, anti-caspase-3, cleaved caspase-3, anti-cytochrome c (1:1000, Bioworld Technology, China). The blots were thoroughly washed with TBST and incubated at 37 °C with the secondary antibody in TBST containing 5% skimmed milk powder for 1 h. After that, the signal was tested by enhanced chemiluminescence (ECL kit, Millipore, USA). Cox-IV was used as an internal reference for proteins in mitochondria, while β-actin was used in cytoplasm. The membranes were imaged and analyzed using the Quantity One Image Analysis Software (Syngene, U.K.) [49].

### Statistics analysis

SPSS 20.0 software was used to analysis the data, and the data was appeared as mean ± SD. Compare the mean using one-way analysis of variance (ANOVA). Several comparative tests were also conducted. In addition, variance homogeneity test was used to test the data. The mean values of homogenous variances were compared by ANOVA and analyzed the differences between the two groups using least significant difference (LSD). If the data did not obey the normal distribution or the variance was uneven, Welch’s t-test was used, and the Games-Howell test was used for further pairwise comparison [50]. Differences were considered significant when *P* <0.05.

## Results

The present study aimed to explore the potential neuronal apoptosis mechanism of hippocampus of depression-like behaviors in CUMS-induced rats by investigating the function of ICA. Through a series of *in vivo* experiments, it was found that ICA ameliorated neuronal apoptosis via inhibiting GR mitochondrial translocation and expression of Bax, thereby preventing the release of cytochrome C into the cytoplasm to activate caspase 3. Therefore, in the data, the function and mechanism of ICA were studied, providing new insights into the pathogenesis of depression.

### ICA ameliorates CUMS-induced depression-like behaviors in rats

First, the body weight was no significant difference among the four groups of rats at the beginning. From 1 to week 5, the body weight in the CUMS group was significantly lower than that of rats in control group from week one to week 5. Specifically, upon the administration of CUMS+ICA and CUMS+Flx, the body weight of the rats increased compared to those of the rats in the CUMS group at the end of the experiment (Figure 2A). On the 35^th^ day, there were significant differences in sucrose preference among the four groups, with the lowest and highest sucrose preference percentages in the CUMS group and the control group, respectively. Compared with the CUMS group, the CUMS+ICA and CUMS+Flx groups showed significantly higher sucrose preference, suggesting that ICA may reduce anhedonia behavior of rats (Figure 2B). EPM and the OFT test were first used to evaluate the locomotor behaviors and anxiety of rats. For OFT test, the CUMS group showed the less time spent and entry frequency in the center zone, while, ICA treatment significantly reversed this phenomenon, and Flx group showed a similar effect (Figure 2C). As shown in Figure 2D, compared with the control group, all the stressed rats showed a significant less frequently and lower time spent in the open arms, while ICA and Flx treatment could increase the time spent and frequency in the open arms compared with CUMS group. As for the FST, the immobility time and frequency of the rats in the CUMS group was significantly increased compared with those of the rats in the control group, while the CUMS+ICA and CUMS+Flx groups reduced the immobility time and frequency induced by CUMS group in rats (Figure 2E). Defecation in the open field is always regarded as an index of the animal anxious state. But in our research, defecation among different groups was only slightly different, and there was no significant difference (Fig.2F). Some studies [51, 52] confirmed that circulating CORT is associated with depression, so we measured the CORT levels in serum of the rats to assess the depression-like state. As shown in Figure 2G, the circulating CORT of CUMS increased significantly, while ICA and Flx could decrease the serum CORT levels.

**Fig. 2.**
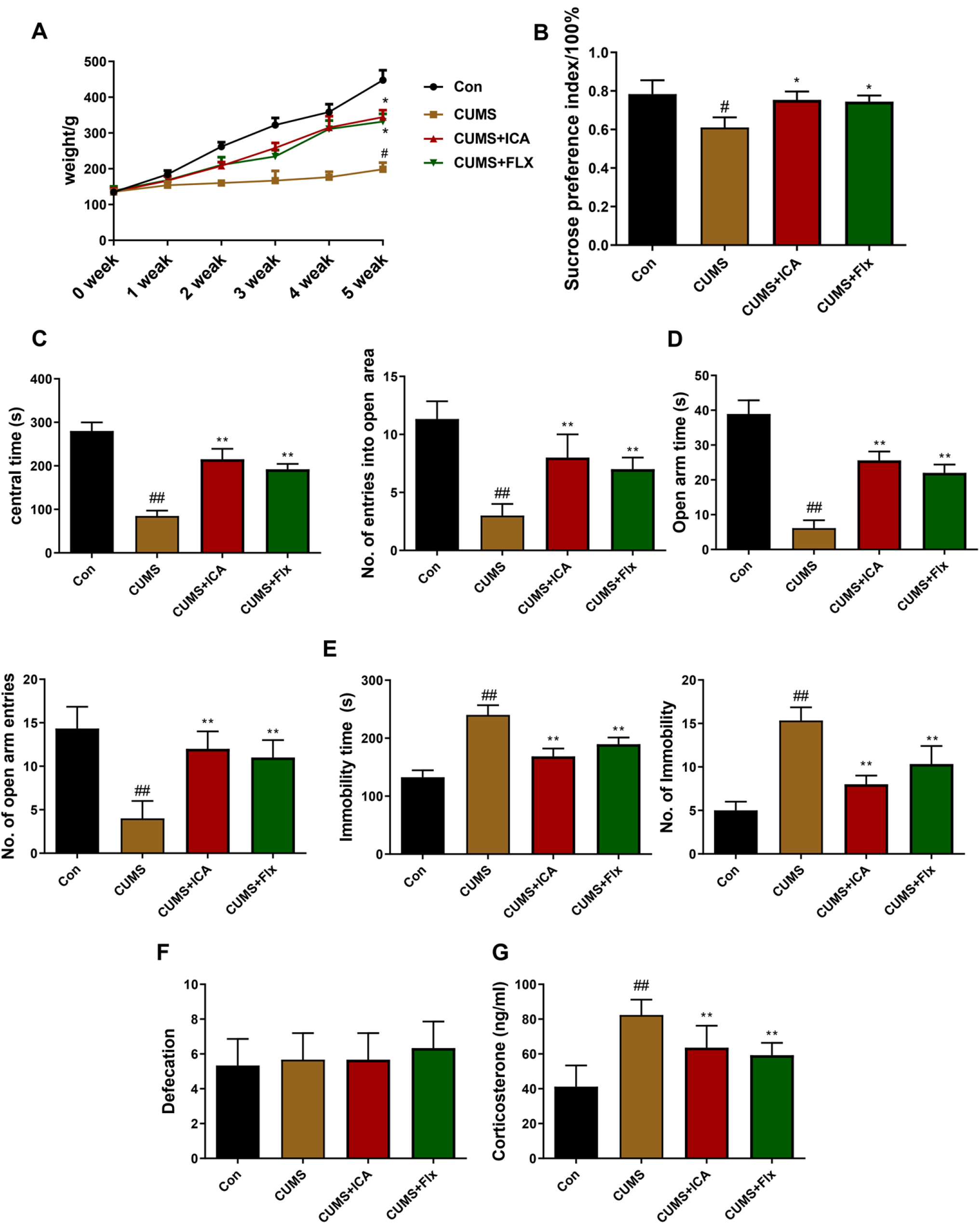
Effects of ICA on the body weight and behavior of rats subjected to different groups. 40 rats were divided into 4 groups (10 rats/group) and treated with Control, CUMS-vehicle (saline 10 ml/kg), CUMS-icariin (20 mg/kg) and CUMS-Flx (positive control, 10 mg/kg) separately for 5 weeks. (A) Changes in body weight from week 0 to week 5. ^#^P < 0.05 vs. Control group, ^*^P < 0.05 vs. CUMS group. df=10. (B) Sucrose preference among study groups at day 35. ^#^P < 0.05 vs. Control group, ^*^P < 0.05 vs. CUMS group. df=11. FST immobile time and frequency. ^##^P < 0.05 vs. Control group, ^**^P < 0.05 vs. CUMS group. df=11. Time and frequency of enter the open arm. ^##^P < 0.05 vs. Control group, ^**^P < 0.05 vs. CUMS group. df=11. (E) OFT center time and frequency were measured. ^##^P < 0.05 vs. Control group, ^**^P < 0.05 vs. CUMS group. df=11. (F) The defecation of rats in open field. (G) The CORT in the serum of rats with different groups. ^##^P < 0.01 vs. Control group; **P < 0.01 vs. CUMS group. df=11. Data were presented as the mean + SD (n = 10 per group).

### ICA decreases neuronal apoptosis in the hippocampal CA1 area of the CUMS rats

Multiple studies suggest that depression induces neuronal apoptosis in the hippocampal CA1subfield [39, 53]. Nischeria is considered a morphological index of the neural cell function [54], while TUNNEL staining is usually used to detect apoptosis [46]. Based on the available evidence, hippocampal neuron morphology changes were observed using Nissl staining. In our results, the normal cell morphology, clear nucleus and purple blue staining of Nissl bodies can be seen in the normal neurons of control, CUMS+ICA and CUMS+Flx groups (Figure 3A, C, D), while the TUNNEL positive cells of CUMS group are shown with apoptotic bodies and pyknotic nuclei (Fig.3F). Taking together, samples from rats chronically administered with ICA showed that neurons in the CA1 subfield of the hippocampi were effectively protected compared to the CUMS exposed rats.

**Fig.3.**
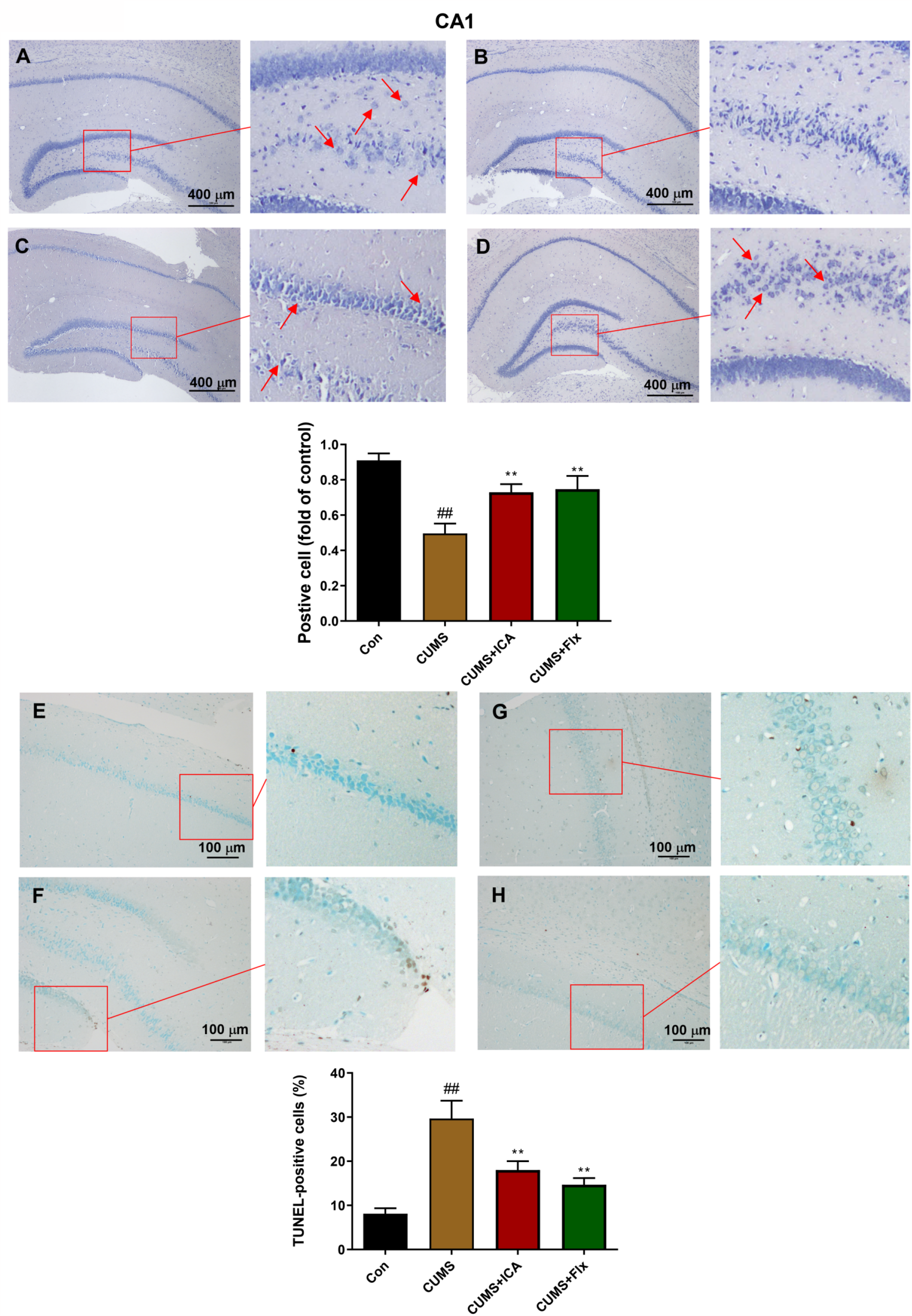
Photomicrographs of hippocampal pathological sections. (A)-(D) Nissl staining of CA1 region in hippocampus. (E)-(F) TUNNEL staining of CA1 region in hippocampus. (A), (E) are Con group; (B), (F) are CUMS group; (C), (G) are ICA group; (D), (H) are Flx group. The red arrows indicate the Nissl bodies and apoptotic bodies. Data were presented as the mean + SD (n = 10 per group). ^##^p < 0.01 vs. Control group; **p < 0.01 vs. CUMS group, df=11.

### ICA prevents mitochondrial translocation of apoptotic proteins

During stress, high concentration GC in serum can induce apoptosis by GR mitochondrial translocation [55]. In the present study, GR and apoptosis-related protein (Bax and Bcl-2) expression in mitochondria and cytoplasmic was detected by western blot (Figure 4A and E). Results showed that GR expression was increased in mitochondria of CUMS exposure group (Figure 4B), while was significantly reduced in cytoplasm (Figure 4F). This effect was significantly subverted by ICA and Flx treatment, indicating that ICA can prevent the translocation of GR from cytoplasm to mitochondria. Similarly, Bax was markedly increased both in the mitochondria and cytoplasm in CUMS group, while ICA and Flx decreased the level of Bax compared with CUMS group (Figure 4C and G). It is well known that the ratio of Bax/Bcl-2 determines the death or survival fate of cell after apoptotic stimulus [56]. In addition, Bcl-2 can regulate MOMP by combination with Bax [57]. In our study, the Bax/Bcl-2 ratio was also analyzed. The data demonstrated that whether in mitochondria or cytoplasm, Bax/Bcl-2 ratio was significantly increased under the stimulation of CUMS compared with the control group, indicating that apoptosis was promoted, and the administration of ICA and Flx decreased the ratio, revealing that the pro-apoptotic ability of CUMS was inhibited (Figure 4D and H).

**Fig.4.**
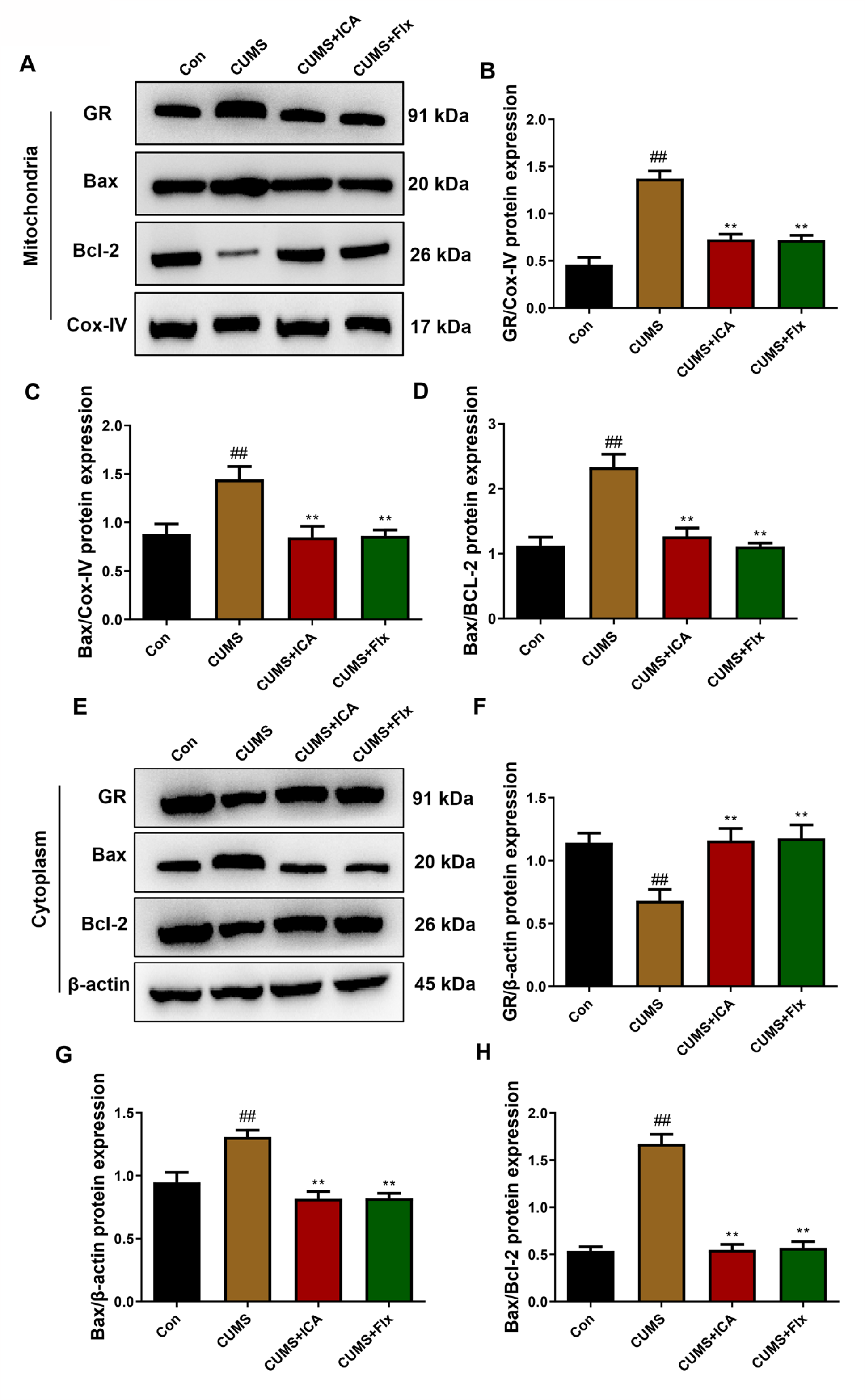
Western blot experiment demonstrating the effect of icariin on mitochondrial and cytoplasmic GR, Bcl-2 and Bax levels in rat hippocampus (A), (F)-(H) icariin treatment inhibited the expression of GR and Bcl-2 of mitochondria in the CUMS hippocampus; (B), (C)-(E) The expression of GR, Bcl-2 and Bax in cytoplasm. Data were presented as the mean + SD. ^##^P < 0.01 vs. Control group; *P < 0.05, **P < 0.01 vs. CUMS group, df=11.

### ICA inhibits caspase activation and cytochrome C release to the cytoplasm in hippocampus

Caspase 3 is the main terminal processing protease and it plays a vital role in cell apoptosis [41]. The release of cytochrome C into the cytoplasm can activate caspase-3, thereby inducing apoptosis [58]. In our study, upon the administration of CUMS, caspase-3 (Figure 5B), cleaved caspase-3 (Figure 5C) and cytochrome C (Figure 5D) expression was increased in the cytoplasm. While under the stimulation of ICA and Flx, the expression of all three proteins decreased compared with the CUMS group. Indicating that the level of cytochrome C extravasation into the cytoplasm is reduced, thereby inhibiting the activation of caspase-3. The above results indicated that ICA can inhibit the MOMP induced by the Bax, thereby preventing the release of cytochrome C into the cytoplasm to cause the activation of caspase-3, and inhibiting the apoptosis of neuronal cells through the Bax/cytochrome C/caspase-3 axis.

**Fig.5.**
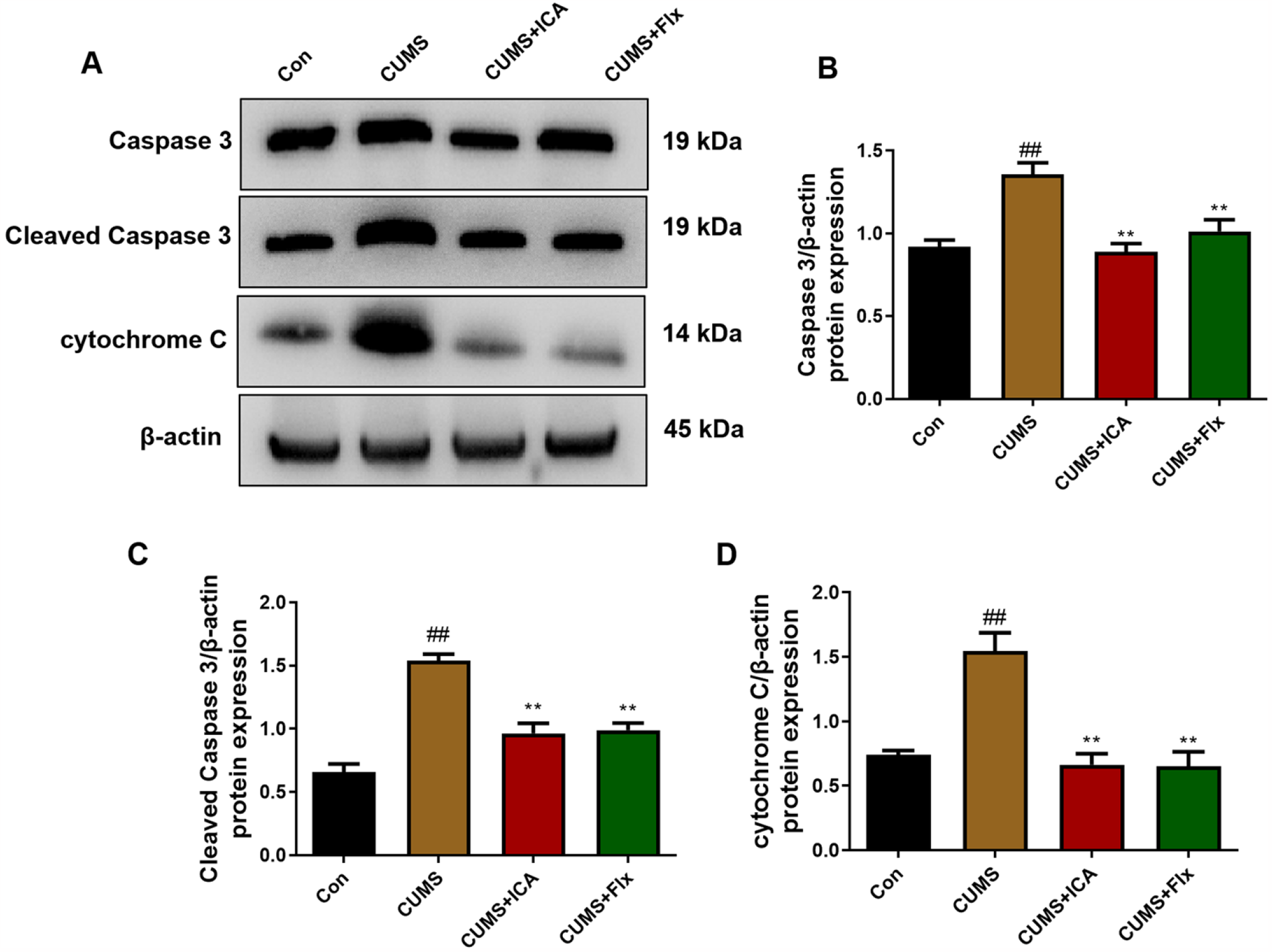
ICA treatment effects on the expression of cytoplasmic apoptotic proteins. (A) Western blot was used to detected the apoptotic proteins (caspase 3 and cytochrome C) in cytoplasmic. The bar graphs reflected the cleaved caspase 3 (B), caspase 3 (C), cytochrome C (D) proteins expression in each group. Data were presented as the mean + SD. ^##^P < 0.01 vs. Control group; *P < 0.05, **P < 0.01 vs. CUMS group, df=11.

## Discussion

ICA is one of the most bio-active compounds purified from the Chinese herb *Epimedium*. Our previous study verifies its broad-spectrum effects on antioxidant [59], anti-tumor [60, 61], anti-inflammation [29] and anti-depression [62]. For example, Zeng et al. have proved that ICA can prevent depression and dysfunctional hippocampal neurogenesis by regulating the certain proteins expression in the cerebrospinal fluid [63]. Otherwise, Liu et al. revealed that ICA exerts antidepressant-like effects on brain tissue by inhibiting of NF-κB signaling activation and enhancing antioxidant status and anti-inflammatory effects by the NLRP3-inflammasome/caspase-1/IL-1β axis [26]. In this study, we analyzed the effect of ICA using the known antidepressant drug Flx as a positive control, and the results documented that five weeks’ administration of ICA prevented the depressant-like behaviors of male rats induced by CUMS and attenuated the hippocampal neuron apoptosis, which consistent with the effect of Flx, and even some effects are better than Flx, such as suppressing the ratio of Bax/Bcl-2. Hence our results support the idea that ICA exerts anti-depressant effects.

GR is a member of the steroid receptor superfamily [66], and the inactive form of GR is related to Hsp-90 in the cytoplasm [67]. In the selection process of GR signal regulated thymocytes, mitochondria can act as an important signal-integrator organelles [18]. Interestingly, GR signaling plays an important regulatory role in hippocampal selection and apoptosis [68]. When GCs bind to their receptor, GR is isolated from Hsp-90 and translocated to the nucleus, where it binds to the target genes and acts as a transcription factor [69]. In physiological conditions, GCs are secreted by the adrenal cortex, regulate the biosynthesis and metabolism of sugar, fat and protein [70], and it also play an important role in the anti-inflammatory process [71]. Conversely, during pathological conditions, GCs combine with the GR, inducing the expression of apoptosis proteins in the cells [72]. The fact that mitochondria directed GR induces apoptosis suggests that the exclusive expression of GR in mitochondria is sufficient to trigger apoptosis [55]. The decrease of mitochondrial GR level detected in males may help to mitigate the adverse effects of LPS on mitochondrial signaling [73]. CORT is also secreted by the adrenal gland upon stress exposure, and multiple studies proved the tight relationship between CORT and depression [74, 75]. Repeated CORT injections in animals result in HPA axis deregulation, cognitive, memory decline and neuronal damage, and induce depression-like behavior [76, 77]. In addition to the well-understood mechanism of CORT, it can also affect mitochondrial functions through binding with the GR within the brain cells and brain tissues [78, 79]. There is evidence that compared with the healthy control group, the GCs level of men who meet the criteria for MDD standard is significantly higher in men compared with healthy controls, whereas no difference is observed in depressed women, and female rats have higher baseline levels of CORT than males [80]. In addition, Brkic et al. revealed that alterations in mitochondrial GR were more prominent in the PFC of males [48]. Therefore, in order to make the results more significant and informative, we choose male rats to establish the animal model. In our study, during exposure to CUMS, in spite of a marked increase in serum CORT, the level of cytoplasmic GR decreased while mitochondrial GR increased. Thus, it seemed that chronic stress caused redistribution of GR by a CORT independent mechanism.

Bax is the main pro-apoptotic protein of the Bcl-2 family which plays an important role in cell apoptosis [81, 82]. Normally, Bax is located in the cytoplasm; it can aggregate into homologous dimers or combine with Bcl-2 to form heterologous dimers [83, 84]. Bax plays an important role in the process of apoptosis as Bax homodimers can bind to the mitochondrial outer membrane, resulting in an increase of MOMP [85, 86]. After Bax formed pores on the mitochondrial outer membrane, cytochrome C is released into the cytoplasm, which is involved in the formation of the apoptosome in conjunction with Apaf1 and caspase-9 and activated caspase-3 [18], after which caspase-3 degrades caspase-activated deoxyribonuclease (CAD). CAD is then released and enters the nucleus to destroy DNA at the nucleosome joint region [87-89]. However, Bcl-2 can regulate mitochondrial membrane permeability by combination with Bax, and then control the release of cytochrome C [90]. Our results show that ICA can reduce the expression of Bax and GR in mitochondria increased by CUMS. In line with this result, the release of cytochrome C into the cytoplasm was inhibited by ICA, further preventing the activation of caspase-3. In summary, our results confirmed that ICA related to its inhibition of neuronal apoptosis in hippocampus through mitochondrial apoptotic pathway. However, some limitations of our study also exist. For example, Luo et al. reported that GR translocation may be reduced under prolonged CUMS stimulation [91]. We have not made a comparison for this, and further research is needed. In addition, we did not detect differences in baseline corticosterone and GR in the sex group, which may indicate that ICA did not produce any sex-specific lasting effect on the neuronal apoptosis. Therefore, in the following work, we will establish a bisexual rat model to investigate whether this apoptotic mechanism is related to sex.

## Conclusion

Taken together, our research provide direct evidence that ICA played an antidepressant role in CUMS rats by decreasing the GR mitochondrial translocation and reducing the neuronal apoptosis in the hippocampus through a mitochondrial apoptotic pathway of Bax/cytoplasm C/caspase-3 axis. Our current findings suggest that ICA may be an effective therapeutic treatment to prevent CUMS-induced depression-like behaviors in rats.

## Disclosure statement

### Ethics approval

All experiments were conducted in line with the guidelines of Animal Care and Use Committee at Affiliated Hospital of Nanjing University of Chinese Medicine(NO:HS-A-2021-0721). Every effort was done to minimize the animals’ suffering.

### Availability of data and materials

The data used to support the findings of this study are available from the corresponding author upon request.

## Acknowledgements

Not applicable.

## Figure Legends

**Fig.S1.**
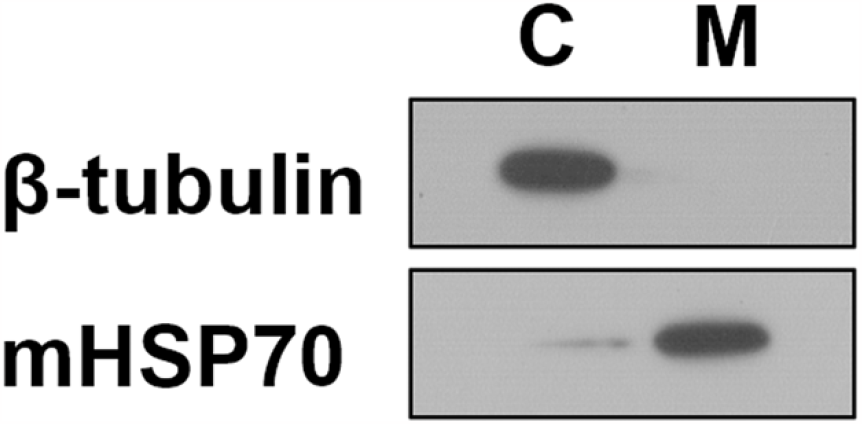
The purity of subcellular fractions. The purity was assayed using specific antibodies against β-tubulin for cytoplasmic and mHsp70 for mitochondrial fraction.

